# Utilizing Protein Bioinformatics to Delve Deeper into Immunopeptidomic Datasets

**DOI:** 10.1101/2024.09.05.611486

**Authors:** Christopher T. Boughter

## Abstract

Immunopeptidomics is a growing subfield of proteomics that has the potential to shed new light on a long-neglected aspect of adaptive immunology: a comprehensive understanding of the peptides presented by major histocompatibility complexes (MHC) to T cells. As the field of immunopeptidomics continues to grow and mature, a parallel expansion in the methods for extracting quantitative features of these peptides is necessary. Currently, massive experimental efforts to isolate a given immunopeptidome are summarized in tables and pie charts, or worse, entirely thrown out in favor of singular peptides of interest. Ideally, an unbiased approach would dive deeper into these large proteomic datasets, identifying sequence-level biochemical signatures inherent to each individual dataset and the given immunological niche. This chapter will outline the steps for a powerful approach to such analysis, utilizing the Automated Immune Molecule Separator (AIMS) software for the characterization of immunopeptidomic datasets. AIMS is a flexible tool for the identification of biophysical signatures in peptidomic datasets, the elucidation of nuanced differences in repertoires collected across tissues or experimental conditions, and the generation of machine learning models for future applications to classification problems. In learning to use AIMS, readers of this chapter will receive a broad introduction to the field of protein bioinformatics and its utility in the analysis of immunopeptidomic datasets and other large-scale immune repertoire datasets.

## 1 Introduction

With the recent success of immunotherapeutics [1–3], there has been a renewed focus on some of the highly diverse molecules central to adaptive immune recognition, specifically T cell receptors (TCRs) and the peptide-major histocompatibility complex (pMHC). Fundamentally, the success or failure of an individual’s adaptive immune system to recognize pathogenic invaders like viruses, bacteria, or cancer hinges on the ability of an MHC molecule to present a pathogenic peptide on a cellular surface, and on the existence of a T cell bearing a receptor which can recognize this pathogenic peptide [4, 5]. While this sounds simple enough in theory, this recognition process is greatly complicated by the fact that the human body can produce on the order of 10^15^, denoted as 𝒪 (10^15^), possible TCR sequences [6], and that the space of possible 12-mer peptides is likewise 𝒪 (10^15^). Further complications arise when we consider that the average human has only 𝒪 (10^12^) circulating T cells, and a given cell only presents 𝒪 (10^5^) peptides at a given time [7]. How then can we parse the signal from the noise and identify the molecules critical for immune recognition in a given immune response?

Recent improvements in next-generation sequencing and mass spectrometry have led to a proliferation of unbiased approaches that are capable of characterizing a large fraction of a given T cell response and a sampling of the peptides presented to these T cells [7]. However, in the absence of functional data (i.e. T cell activation or peptide immunogenicity measurements) how can we consistently identify those TCRs or peptides that are critical for initiating an immune response? Absent restrictions in resources and time, we would purify and test every sequenced TCR-pMHC pair and identify those that show strong binding and responses to pathogenic peptides that are commonly expressed on cell surfaces. Given the aforementioned magnitude of these repertoires, and our collective inability to allocate endless resources or time, this is experimentally infeasible. This necessitates a computational approach to processing these large sequence datasets. While structural prediction has advanced greatly through breakthroughs such as AlphaFold [8], even the best software still struggle to predict TCR-pMHC complex structures. This, finally, leads us to the goal of this chapter, which is the use of protein bioinformatics to systematically characterize large sequence datasets.

The Automated Immune Molecule Separator (AIMS) was created solely for such protein bioinformatic analysis of large sequence datasets [9]. AIMS takes a pseudo-structural approach to bioinformatic analysis, avoiding making explicit structural predictions while maintaining relative structural context. For instance, in AIMS analyses, the 6 CDR loops of antibodies (Abs) and TCRs are encoded as distinct entities. Likewise, the anchor regions and TCR-contacting residues of peptides are typically encoded as distinct features. Maintaining this structural context and position-sensitivity allows users to isolate binding hotspots and key features in each individual repertoire. Whereas explicit structural prediction provides increased precision (TCR residue X interacts with peptide residue Y), this comes with low accuracy, which can lead to the generation of incorrect hypotheses. AIMS, conversely, reduces the precision of the analysis with the benefit of increased accuracy (TCR region A shows strong biophysical compatibility with peptide region B).

While AIMS can be used for interaction prediction and the analysis of TCRs, MHC, peptides, antibodies (Abs), and even non-immune molecules [See **Note 1**], we will here focus on the analysis of immunopeptidomic datasets. This chapter will outline the best practices for installing and using AIMS, and walk readers through the key features of AIMS, starting with the initial pseudo-structural encoding, moving through the position-sensitive biophysical property analysis and statistical sequence feature quantification, and ending with the creation of machine learning models from acquired datasets.

Throughout this chapter, there will be references to **Notes**, which have additional information that would clutter the otherwise linear protocol outlined below. AIMS is, however, inherently a nonlinear analysis tool. Re-analysis of data with certain settings and options changed is strongly encouraged. The **Notes** provided in **Section 4** will provide helpful hints for how to do this, along with other more detailed explanations of potentially confusing steps.

## 2 Materials

The AIMS software can be installed and run on a standard desktop or laptop (AIMS was developed and tested on a 2.3GHz, 8-Core Intel i9 laptop with 32GB RAM) or alternatively can be deployed on a high-performance computing cluster. All functions exclusively utilize CPUs, with optionality for parallelization of code and some support for GPU calculations being added as the software continues to evolve. The software is operating system independent, although Windows OS users may need to take a few extra steps to run the software compared to Mac or Linux OS users [see **Note 2**]. While AIMS can be installed with a oneline command, this section will take users through the “best practices” for safely installing AIMS and creating stable, isolated programming environments using Conda (see **Note 3** for an explanation of why Conda is important).

### 2.1 Create an Isolated Programming Environment Using Conda

1. Install Anaconda (www.anaconda.com/products/individual) or miniconda (docs.conda.io/en/latest/miniconda.html) [see **Note 2** for distinction]
2. Open a terminal (Mac or Linux OS) or console window (Windows OS, see **Note 2**). All following steps will include code entered into the terminal or console window
3. Create a new Conda environment by entering: `conda create -n aims-env python=3.7`
4. If Anaconda/Miniconda is installed properly, a Y/N prompt should appear. Enter `y` to create a Conda environment
5. Test for proper environment creation by activating the environment: `conda activate aims-env`
6. You should now see extra text in your terminal command line that reads `(aims-env)`
7. Your Conda environment is now activated, and you have a stable space to install AIMS. When you would like to exit the environment, enter: `conda deactivate`

### 2.2 Installing AIMS

1. Ensure that the AIMS Conda environment is activated (see **Section 2.1, Step 5**)
2. Install the Python package manager pip using: `conda install pip`
3. Install AIMS with one command: `pip install aims-immune` pip will automatically install every package needed for proper functionality of AIMS
4. AIMS is now installed, and ready for use. If needed, see **Note 4** for installation troubleshooting and alternative installation options
5. Figure 1. provides a visual example of all **Section 2** steps entered into the terminal

**Figure 1:**
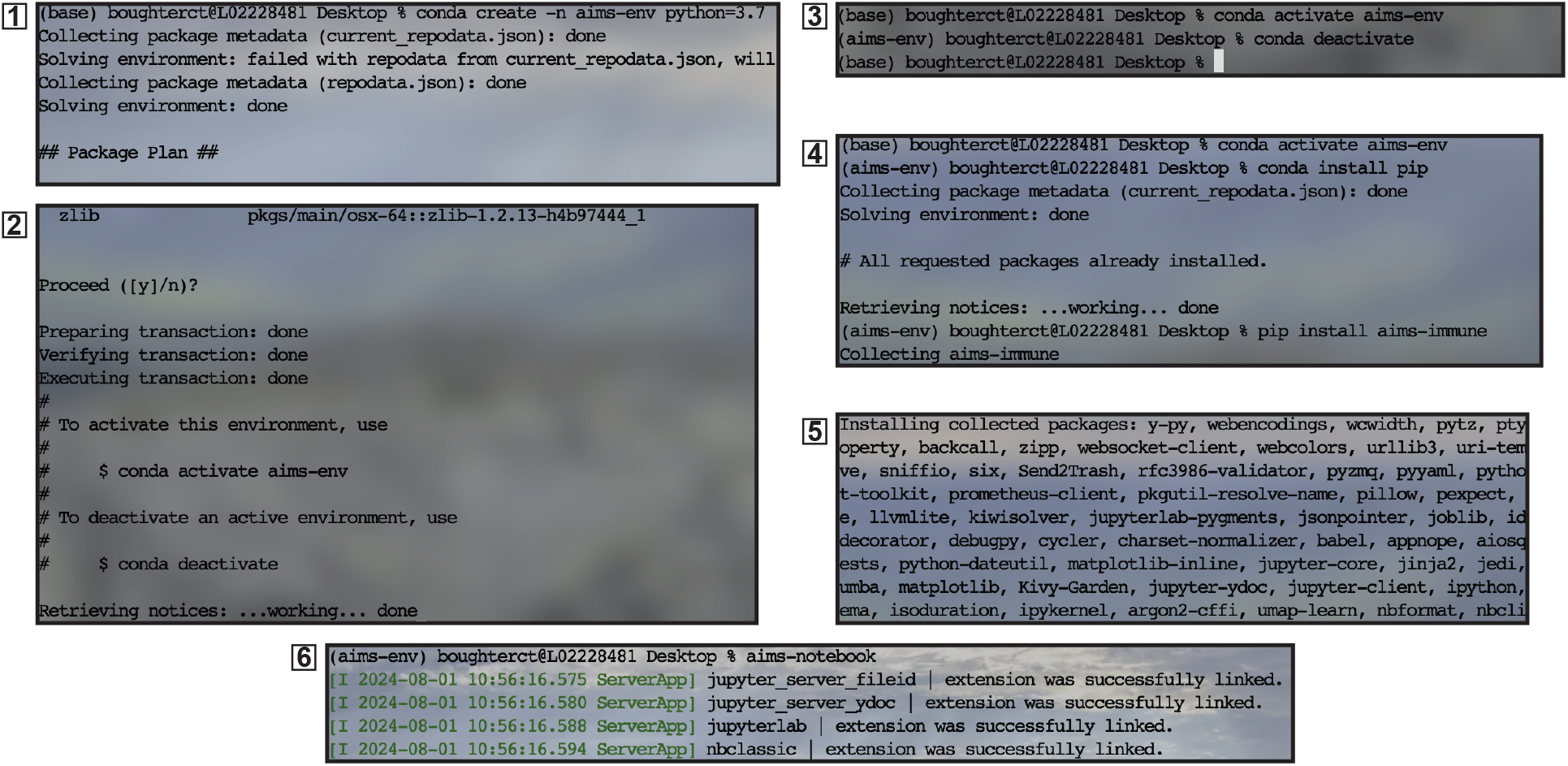
Screenshots taken directly from the terminal in the process of: creating a Conda environment (1), environment installation (2), environment activation and deactivation (3), AIMS download (4) and installation (5), and finally launching of AIMS (6). Most screenshots are only partial images of the full installation process. User input here is denoted by the presence of “%”.

## 3 Methods

Once AIMS is installed and users have their data properly formatted (see **Note 5** for details on formatting), they can initiate data analysis. AIMS users are encouraged to re-run the analysis using combinations of different analytical approaches provided within this chapter to uncover the key features of their data. Additionally, AIMS can be run in three distinct modes (see **Note 6**). For clarity, here the analysis pipeline will be discussed in a linear fashion and discuss each analytical tool in the context of a Jupyter Notebook (see **Notes 6 and 7**).

### 3.1 Starting the Software

*[Optional]* To follow along with the example data used in this chapter, copy test data to the current directory by entering in the terminal: aims-tests. A folder called “test data” will appear in your current directory, containing the relevant input files.

1. If you haven’t already, follow the steps in **Section 2** to ensure you have installed Conda, AIMS, and have a working Conda environment from which to run AIMS
2. Open the terminal or console application (see **Note 2**)
3. Activate (or re-activate) your Conda environment using: `conda activate env-name`
4. Start AIMS (one of the three modalities) by entering: `aims-notebook` See **Note 7** for an important trick when using the Jupyter notebooks
5. All subsequent steps involve changes to, or execution of, code within the newly opened Jupyter notebook. There are no more commands entered into the terminal or console

### 3.2 Load Data & Define Key Analysis Features

In this step, we will load in data and define where data should be saved. In this chapter, all example data will come from MHC Class I-derived peptides identified in the HLA Ligand Atlas [10] isolated from kidneys and pancreas in multiple human donors.

1. Define the directory where the data is located (see **Note 8** on navigating directories): `datDir = ‘ ∼/Desktop/test_data/peptides/ ‘`
2. Define the directory/folder where outputs should be saved to (see **Note 8**): `outputDir = ‘ ∼/Desktop/aims_analysis ‘`
3. Specify the filename(s) of the data being used in the analysis: `fileName=[‘hla_atlas_kidney.csv’,‘hla_atlas_pancreas.csv’]`
4. Specify the names given to these datasets that will be used in generated figures: `datName = [‘ Kidney’, ‘Pancreas’]`
5. Specify additional options for the analysis, for instance removing duplicate entries from each dataset using: `drop_duplicates = True` See **Note 9** for a thorough list of analysis options that can be defined from the outset of AIMS initialization

### 3.3 Encode Sequences into a Biophysical Property Matrix

This is the final step primarily focused on generating data that will be later analyzed elsewhere in the pipeline. This is one of the more computationally intensive steps, so be sure to limit the use of other memory-intensive programs.

1. Specify the alignment scheme, either center, left, right, or “bulge” (see **Note 10**). If choosing “bulge” specify the bulge pad: `align=‘bulge’; pad = 6`
2. Generate the encoding based upon this alignment specification. This will produce the plot seen in **Figure 2**. Each amino acid is represented by a given number, while gaps in the alignment are represented by a “0”, or white space in the figure
3. Specify the normalization to be used for the amino acid biophysical properties. To rescale all 62 biophysical properties (see **Note 11**) to a norm of 1, set: `normalize = True` To re-normalize these properties based on their sequence diversity (see **Note 12**), i.e. prioritize the most diverse regions of the peptide dataset, set: `renormalize= True`
4. Use the initial encoding to create a high-dimensional biophysical property matrix. AIMS automatically generates this matrix with the additional features specified in the previous steps

**Figure 2:**
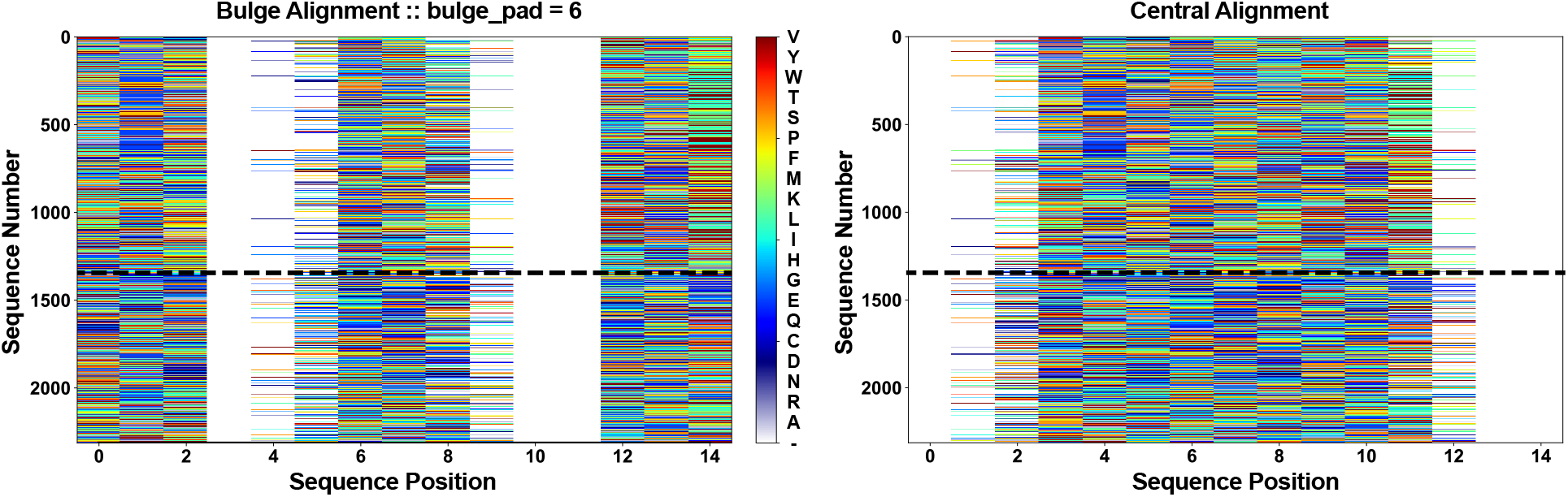
Comparison of the two major types of AIMS sequence encoding. (Left) The bulge alignment, with bulge pad = 6. (Right) The central alignment, generated using the same data as in the left panel. Dotted line separates peptides derived from kidney (top) or pancreas (bottom). Each amino acid is represented by a unique color, while gaps in the alignment are represented by white space.

### 3.4 Identify Biophysically Distinct Clusters of Sequences

In this step, the high-dimensional biophysical property matrix generated in **Section 3.3** is projected into a lower dimensional space for visualization of patterns and clustering of biophysically similar features. This is perhaps the most open-ended of all analysis steps, and one in which users are encouraged to test multiple projection and clustering algorithms.

1. Determine which version of the biophysical property matrix should be used for projection. In most applications, it is recommended that `dchoice = ‘parse’` (see **Note 13** for possible reasons to use other options)
2. Choose the dimensionality reduction algorithm to use. The choice of dimensionality reduction algorithm strongly effects how the sequences are clustered (see **Note 14**). Typically, UMAP is recommended: `reduce=‘UMAP’`
3. Select the clustering algorithm to identify biophysically similar sequences in the projected space. Again, the choice of clustering algorithm is dependent on the structure of the input data (see **Note 15**). Setting `clust = ‘dbscan’` can typically provide reliable clustering for most datasets
4. Set cluster algorithm-specific parameters as needed (see **Note 16**). Users may need to come back to this step multiple times to change parameters `(min_samples, eps, min_cluster_size`, etc)
5. (Optional) Incorporate metadata into sequence cluster plots (see **Note 17**)
6. Plot clustered sequences. Users have an option to display 2D or 3D projections, and overlay either cluster information or metadata information, or both (**Figure 3**)
7. Quantify cluster purity based upon the metadata associated with each sequence. Users can additionally quantify the statistical significance of enrichment of a given population within a cluster using a bootstrapping procedure (**Figure 4**)

**Figure 3:**
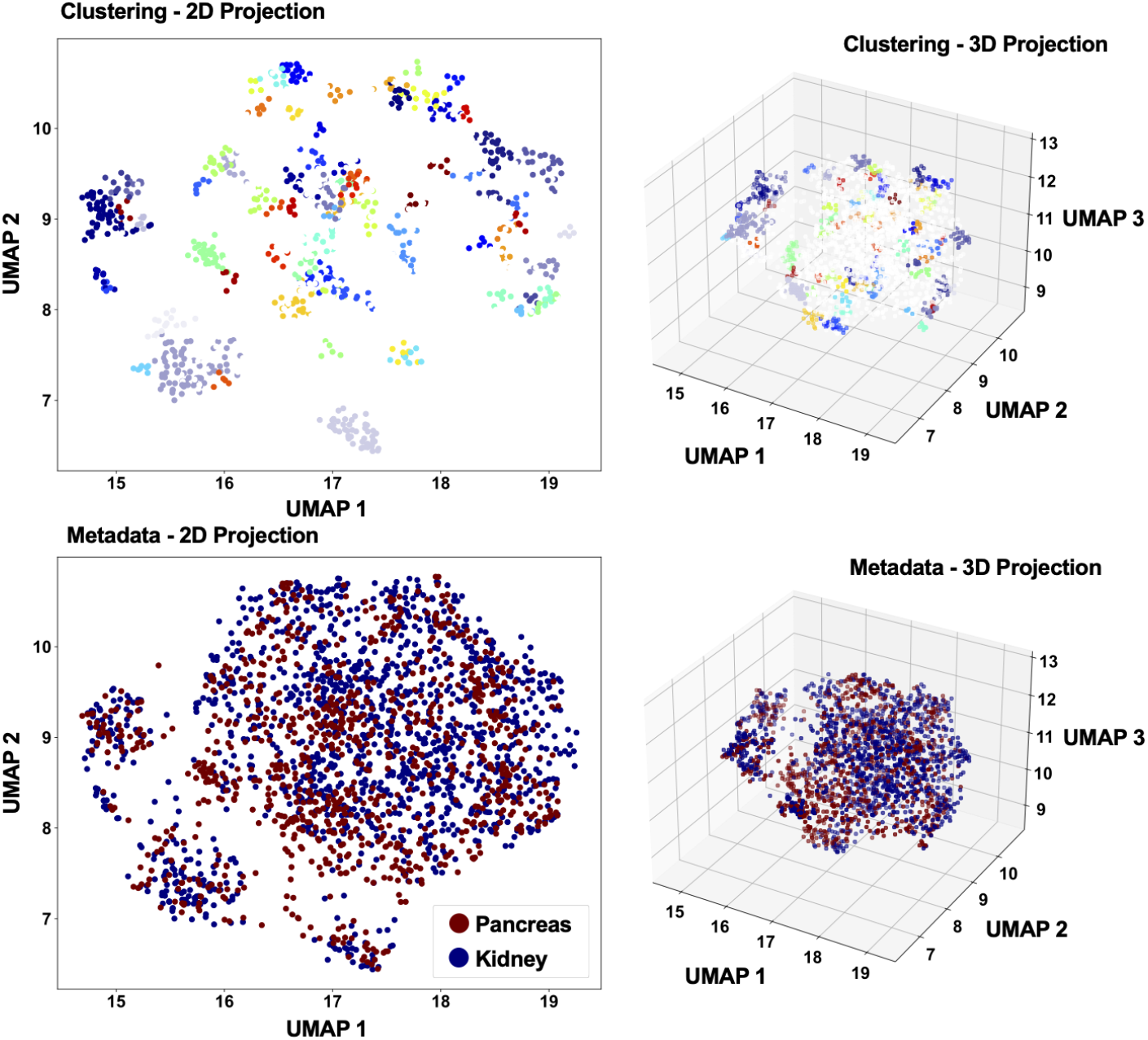
Expected outputs from the AIMS dimensionality reduction and clustering steps. Each individual point is a sequence embedded into UMAP space. (Top) Each sequence is colored based on cluster membership, with each unique color representing a unique cluster. Unclustered sequences are colored white. (Bottom) Each sequence is colored based on sequence metadata, here the tissue of origin. Data is projected either in two UMAP dimensions (left) or three (right). Clustering algorithm: DBSCAN with eps = 0.15.

**Figure 4:**
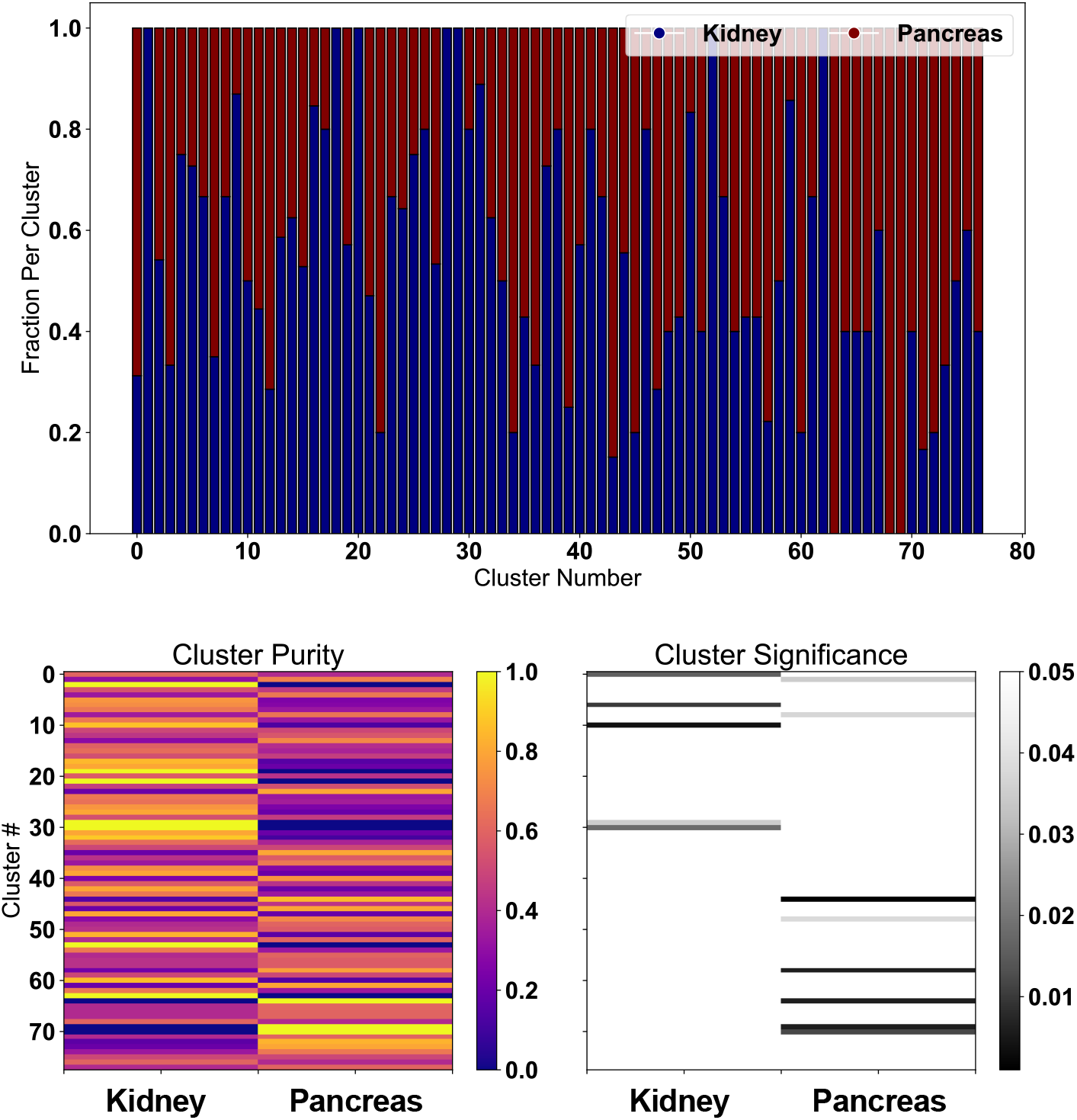
Each cluster identified by the dimensionality reduction and clustering algorithm (here using the data from **Figure 3**) has cluster purity quantified as a fraction of the sequences in each cluster (top). Clusters composed of exclusively kidney-derived peptides are blue, while those comprised only of pancreas-derived peptides are red. This purity can be re-visualized as a heat map (bottom, left) and assessed as a statistically significant enrichment over background (bottom, right). p-values are reported in the color bar, with *p >* 0.05 colored white.

### 3.5 Directly Compare Biophysical Properties of Selected Populations

Next, we select the data we want to further biophysically characterize using the AIMS position-sensitive sequence analysis. See **Note 18** for a discussion on best practices for subset selection and re-analysis of single datasets.

1. If you’d like to analyze distinct clusters of data, set `subset_sel = ‘cluster’` otherwise, set `subset_sel = ‘metadata’`
2. Visualize the encoded sequences re-ordered based upon the subset chosen in **Step 1**. By default, every cluster or unique metadata entry will be visualized in this matrix. See **Note 19** for visualization options for this step and **Figure 5** for example outputs
3. (Optional) Quantify sequence distance between peptides in each cluster. Distance is a useful metric for inter- and intra-cluster or metadata sequence similarity. Inspired by TCRdist [11], the AIMS sequence distance is a simple Euclidean distance in the high-dimensional biophysical property matrix
4. Specifically isolate a subset of the clusters visualized in **Step 2**, and re-plot them (**Figure 6**). Depending on the clusters or metadata chosen, you should see an enrichment of specific motifs. Analysis is recommended for two clusters or metadata subsets, but more can be analyzed simultaneously if needed
5. Plot position-sensitive biophysical property matrices of each of the data subsets (**Figure 7A, B**). These matrices can then be averaged across each sequence in each subset to create a position-sensitive average (**Figure 7C, D**) or averaged across both axes to create a more traditional sequence average for biophysical properties (**Figure 8**)
6. Calculate the statistical significance of the metrics reported in this section (See **Note 20** for details on statistical significance) using a bootstrapping procedure (See **Note 21** on the choice of bootstrapping for significance testing)

**Figure 5:**
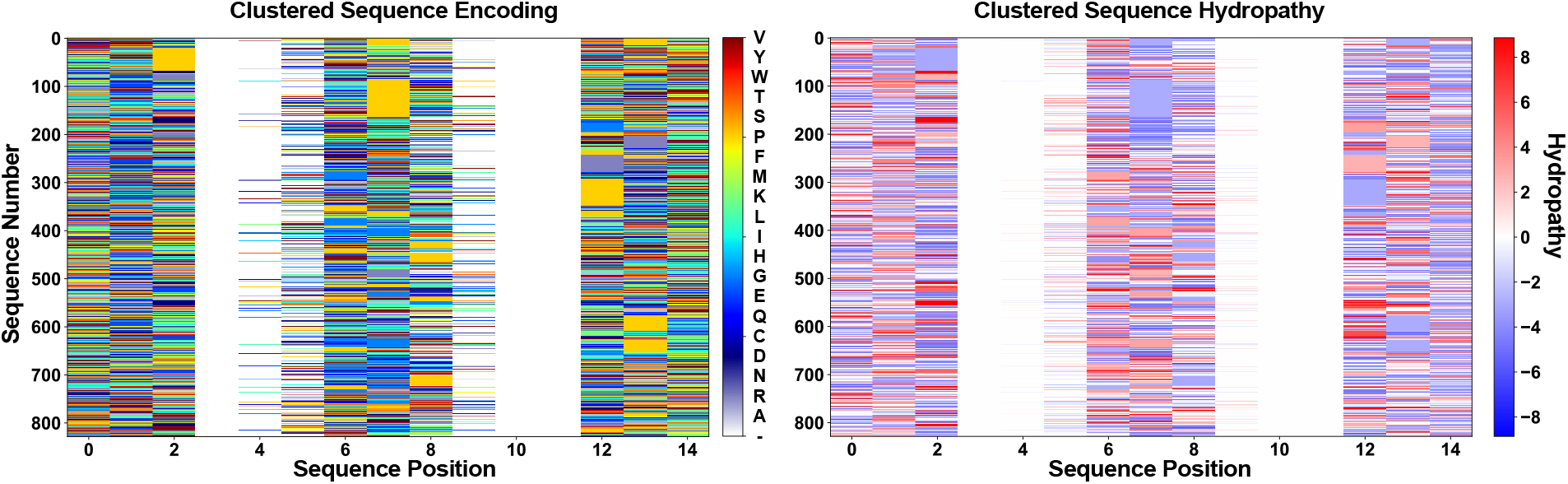
After dimensionality reduction and clustering, sequences can be re-visualized in the encoded AIMS matrix re-ordered by cluster membership. Users can choose to visualize these sequences based on the original amino acid encodings (left) or biophysical properties of these sequences (right).

**Figure 6:**
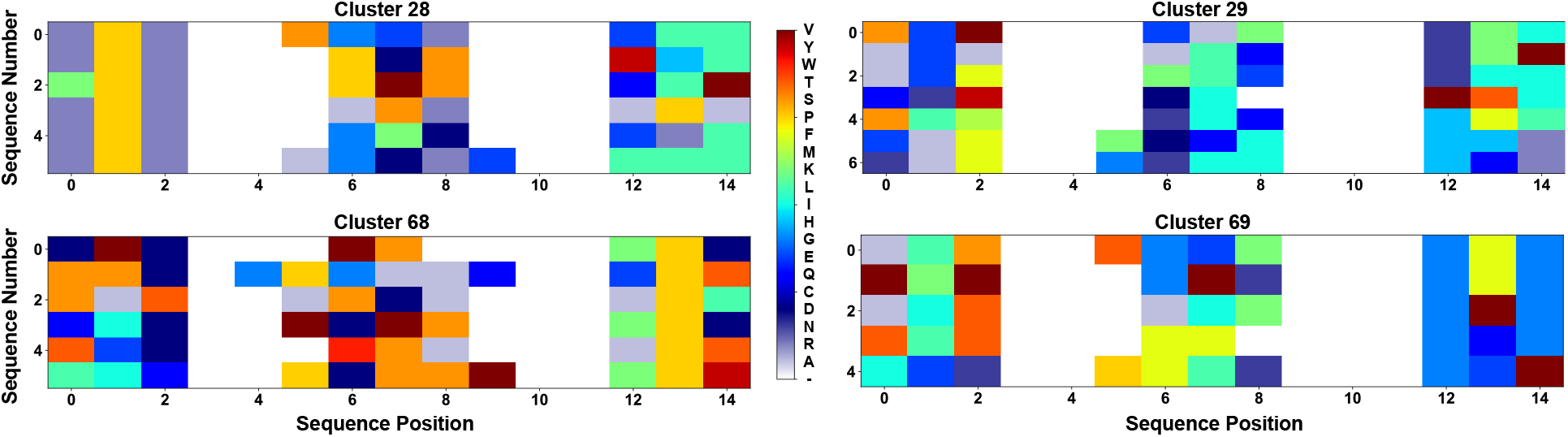
Example figures generated by selecting individual isolated clusters from the re-visualized matrix in **Figure 5**. Here 4 distinct small clusters are isolated for further characterization. Clear enrichments in conserved sequence features are evident from solid vertical lines of color in each panel.

**Figure 7:**
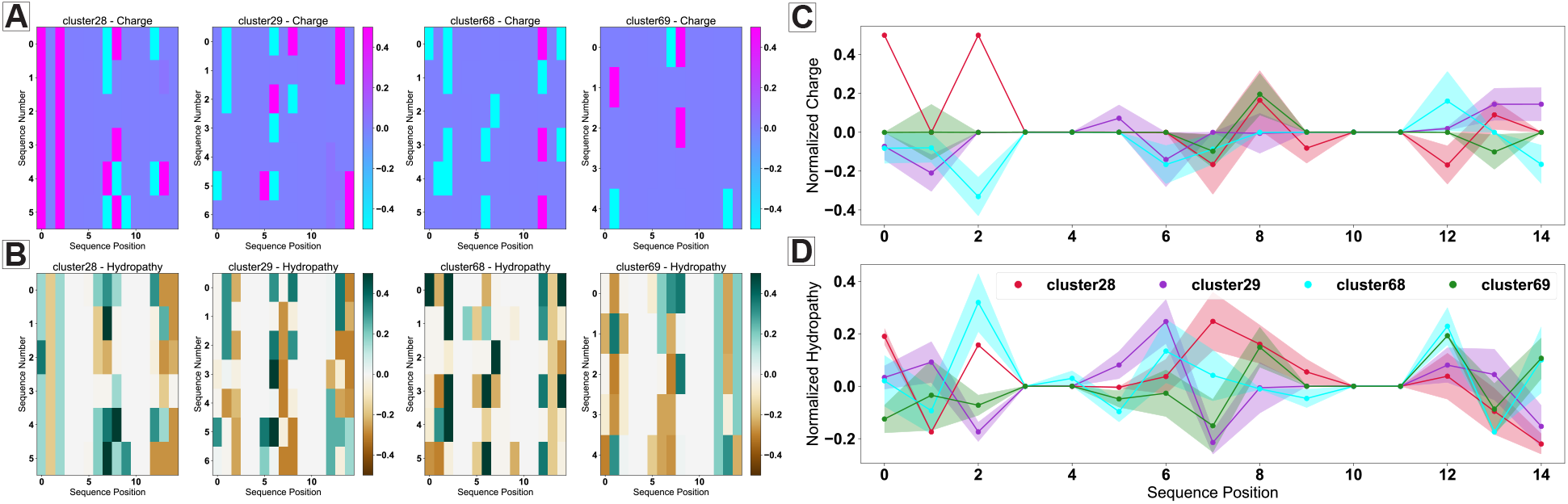
The individual clusters isolated in **Figure 6** are then each characterized using position-sensitive biophysical property analysis. Each cluster can be re-visualized on a per-sequence basis for individual biophysical properties (A, B) or as per-cluster averages that maintain position-sensitivity (C, D). Here charge (A, C) and hydropathy (B, D) are featured, but any of the 62 biophysical properties can be visualized in these steps. In position-sensitive property averages, solid lines give bootstrapped averages while shaded lines give bootstrapped standard deviation.

**Figure 8:**
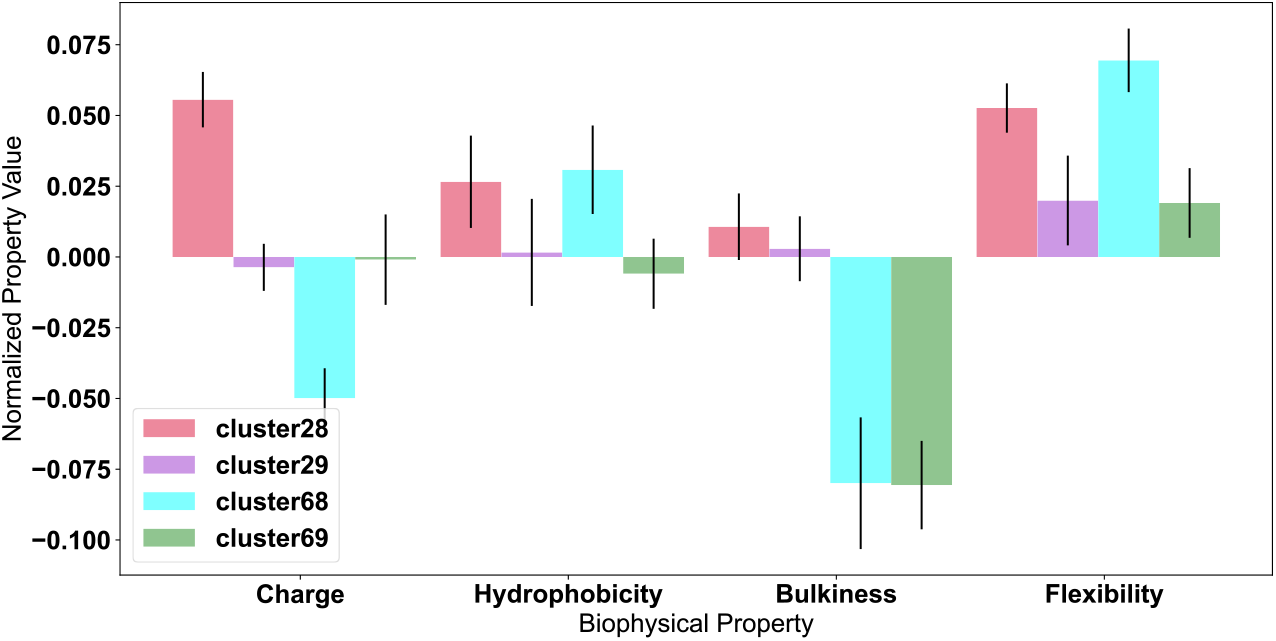
A more traditional per-cluster net average of individual biophysical properties can also be generated from the matrices in **Figure 7**. Values are normalized as per **Note 11**, and averages and standard deviations are generated via a bootstrapping procedure discussed in **Note 21**.

### 3.6 Utilize Information Theory to Delve Deeper into Datasets

After completing clustering and biophysical property analysis for specific subsets of larger datasets, Information Theory can be used as an all-encompassing analysis to identify broader statistical trends in sequence repertoires. See **Note 22** for a primer on Information Theory and the mathematical equations that underlie these metrics.

1. Calculate the sequence coverage for each dataset, identifying regions where direct comparisons of position-sensitive entropy are invalid (**Figure 9A**)
2. Calculate the Shannon entropy of the chosen datasets (**Figure 9A**). Optionally calculate statistics and bootstrapped standard deviation of these metrics (See **Note 23** for best practices specifically for Information Theory statistics)
3. From this Shannon entropy, the mutual information is then calculated (**Figure 9B**). Mutual information identifies strongly coupled sequence positions across each dataset. See **Note 24** for a more thorough discussion of interpretation of mutual information
4. Dissect the Shannon entropy and mutual information quantifications by plotting the position-sensitive amino acid frequencies for each dataset (**Figure 9C**)
5. Calculate the N-gram frequencies of each dataset. Typically, di-gram frequencies are used for ease of visualization (**Figure 9D**), but higher order calculations can be carried out as well (**Note 25**)

**Figure 9:**
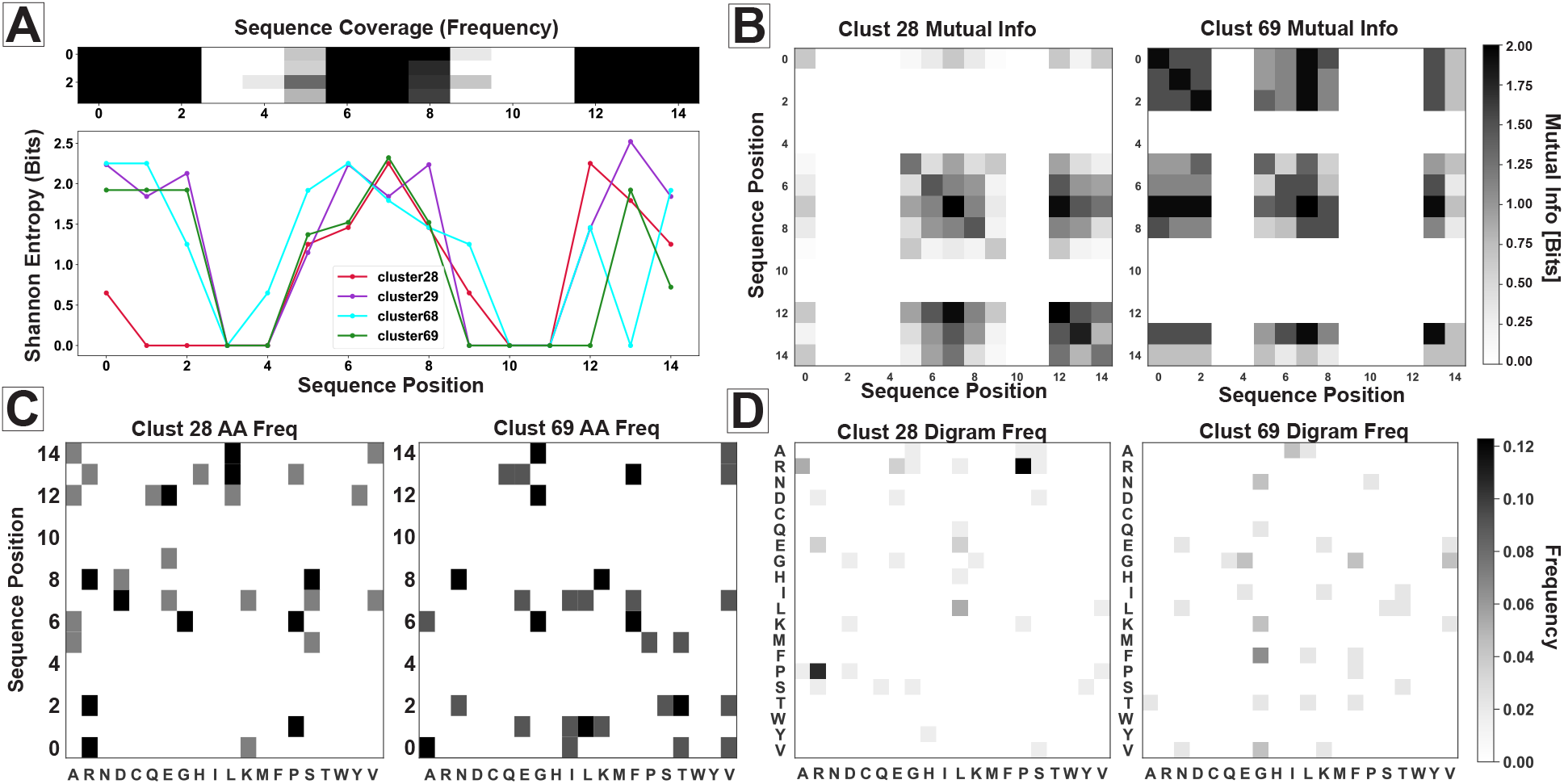
Example outputs of information-theoretic metrics and statistical patterns in input sequences. (A) Sequence coverage (top) and Shannon entropy (bottom) of the sequences isolated from the clusters of **Figure 6**. (B) Mutual information calculated between each position in the sequences in cluster 28 and cluster 69. Position-sensitive amino acid frequencies (C) and digram frequencies (D), provide additional context for the results in (A, B).

### 3.7 Binary Comparisons of Statistical Sequence Features

After quantifying the Shannon entropy, mutual information, and amino acid frequencies of each dataset, direct comparisons by simple differences help to highlight striking features and trends. This necessitates a binary comparison.

1. Visualize the differences in both the mutual information (**Figure 10A**) and optionally calculate a position-sensitive statistical significance testing (see **Note 23** on significance testing for information-theoretic metrics). See **Figure 10B** for an example of a position-sensitive significance testing plot for mutual information
2. Generate a figure for the position-sensitive amino acid frequency differences (**Figure 10C**) and optionally calculate position-sensitive statistical significance
3. Plot the amino acid di-gram difference between the two comparison datasets (**Figure 10D**) and optionally the statistical significance of these differences

**Figure 10:**
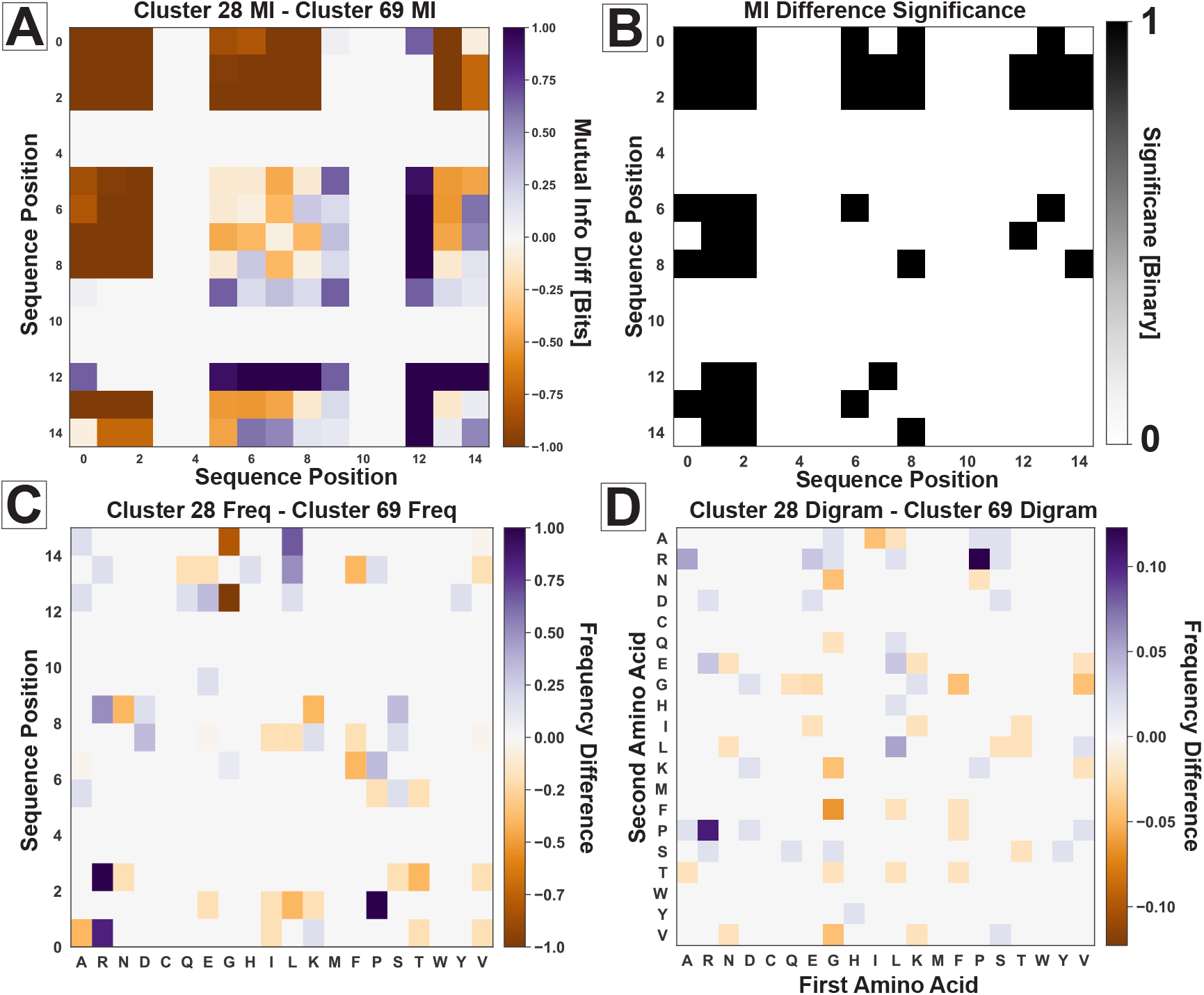
Binary differences in the metrics quantified in **Figure 9** provide key insights into the striking differences between these two clusters. Differences in the mutual information (A) and the statistical significance of these differences (B, *p <* 0.05) provide quantification of sequence cross-talk. Likewise, differences in the position-sensitive amino acid frequency (C) and the amino acid digram frequency (D) highlight strong biases in amino acid usage patterns. Purple colors suggest a bias in these metrics towards cluster 28, while orange colors suggest a bias towards cluster 69. Significance in Panel B is given as a binary, colored black if *p <* 0.05.

### 3.8 Generate Classification Algorithms from Large Datasets

Once the sequences in input datasets have been comprehensively characterized, users have the option of generating machine learning models from their data. This again must be a binary comparison of two datasets (e.g. from two tissues, two treatments, etc.). We here outline the steps for a Linear Discriminant Analysis (LDA)-based model, but other models can be trained using AIMS (see **Note 26**).

1. First, select the datasets which will be used for training the binary classification algorithm. See **Note 27** for some considerations when building a machine learning model
2. Determine the appropriate parameters (typically referred to as hyperparameters in machine learning) for the datasets being used for training. These include `matSize, ridcorr, and pca_split`. See **Note 28** for hyperparameter discussion
3. Run the LDA calculation and visualize the scored sequences (**Figure 11A**). The accuracy of the classification is reported within the Notebook
4. Visualize the top N weights used to generate this classification (**Figure 11B**). The default is `show_top = 5`, but can be adjusted to visualize more weights

**Figure 11:**
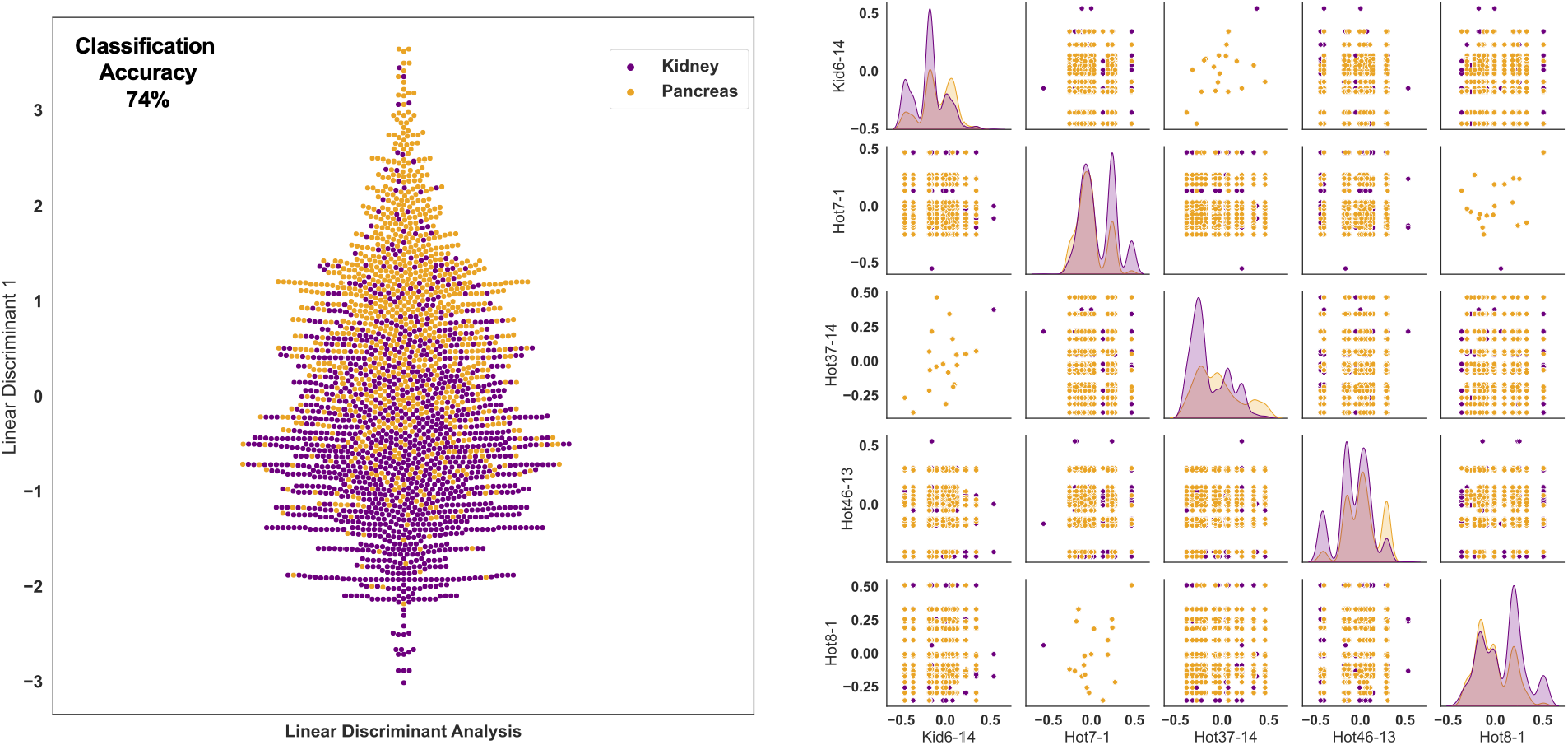
Machine learning algorithms can be used to create classifications of input data, here using the full input dataset. (Left) Sequences projected onto a trained linear discriminant provide an implied likelihood that each sequence is isolated from the kidney (negative values, purple dots) or from the pancreas (positive values, orange dots). Classification accuracy is provided in the figure. (Right) The top five discriminating features are visualized as either histograms (diagonal) or as scatter plots of two features (off-diagonal). Each column or row is labelled by the feature of interest (e.g. Kid6-14 means that Kidera factor 6 at position 14 is a key discriminating feature.

## 4 Notes

- **Note 1:** While this chapter focuses on the use of AIMS specifically for immunopeptidomics analysis, it is more broadly a versatile tool for the analysis of all protein sequence datasets. AIMS has been applied to the analysis of antibodies [12], TCR-pMHC interactions [13], SARS-CoV-2 spike proteins [14], and even to multi-sequence alignments of non-immune proteins [15]. Readers are encouraged to look to these manuscripts for inspiration and a survey of the depth of analysis possible in AIMS.
- **Note 2:** Mac and Linux OS both have Linux-based terminals (In MacOS, simply the “Terminal” app) that make launching Python-based programs simple. Things are a bit more complicated in Windows OS, requiring additional programs to initiate and install AIMS. The full version of Anaconda can provide this, allowing Windows users to both manage environments via the Anaconda GUI as well as execute code using applications such as Qt Console, Spyder, or JupyterLab. However, Anaconda requires nearly 1GB of disk space for download, so MiniConda provides only the essentials for using Conda environments in Terminal for MacOS and Linux users, resulting in a much smaller download size.
- **Note 3:** While users can simply use pip to install AIMS directly, it is best practice to install new software and projects in a self-contained environment, in this case Conda environments. The Python package Kivy, which is used to run the AIMS GUI, tends to cause issues when installing other Python packages. The self-contained AIMS environment can help alleviate this issue, and can prevent software clashes if other packages are installed.
- **Note 4:** If following the installation exactly, users should not run into issues. However, if errors arise while following these steps, the common user errors include: not having the Conda environment activated (**Section 2.1, Step 5**) during either installation or AIMS initiation, having the wrong Python version installed in the environment (not specifying python=3.7 in **Section 2.1, Step 3**), or operating-system based issues. If issues persist, more advanced users can directly download and install AIMS via GitHub: github.com/ctboughter/AIMS
- **Note 5:** Formatting of inputs in AIMS varies depending on the data type being used. Only two types of input formats are used in AIMS: either a comma separate values file (*.csv) or a FASTA-formatted file (* .fasta). Immunopeptidomics, antibody, TCR, and metadata inputs are *.csv formatted, while MHC or MSA analyses utilize FASTA inputs. If data is loaded into Microsoft excel or similar programs, there is a simple setting to save the data as a CSV.
- **Note 6:** This chapter has been written in a way that should apply to all implementations of AIMS (either `aims-cli, aims-gui, or aims-notebook`). However, it should be noted that due to limited resources available for development of AIMS, typically the Jupyter notebook version of AIMS (`aims-notebook`) is the most up-to-date with the most flexibility of features. Users are encouraged to run all analysis in the notebook, if comfortable.
- **Note 7:** The `aims-notebook` command entered in the terminal should automatically launch a new Jupyter Notebook. The notebook is created in such a way that users should only need to execute each cell, one by one, in order, with only small changes made to specify file names and output directories. Cells are executed by either clicking the appropriate button in the window or highlighting each individual cell by clicking on it and hitting (`control+enter` on MacOS) on the keyboard. Lastly, if custom changes are made to the notebook that users would like to save, they need to first save the notebook (`command+s` on MacOS) and then next time they go to start the notebook, type `jupyter lab` in the terminal, rather than `aims-notebook`. Each instance of `aims-notebook` creates a new notebook with a date in the name, whereas the command `jupyter lab` simply opens the Jupyterlab app where previously generated notebooks can be initiated.
- **Note 8:** For novice programmers, every folder and file on your computer is associated with a path to it. Think of this as a list of the folders you typically click through. If you are running your notebook from within a folder called “`analysis`” and keep your data in a folder called “`data`” within this analysis folder, then the path to that folder is `analysis/data`. A file in the `data` directory “`sequences.csv`” then has the path `analysis/ data/sequences.csv` . Lastly, in naming files for input and output in AIMS, it is recommended to replace any spaces with a dash or underscore (i.e. change “`my_peptides.csv`” to “`my_peptides.csv`”. Spaces in folder or file names can complicate directory navigation.
- **Note 9:** Early AIMS user options allow for alterations in the analysis in the following ways:

- `subset = [True/False]`: not applicable to peptide analysis. Allows users to take slices of FASTA-formatted sequences, which are typically 100 amino acids or more
- `num_loop = [Integer (i.e. 1, 3, 6)]`: not applicable to peptide analysis. Defines the number of loops in antibody or TCR analysis to use
- `drop_duplicates = [True/False]`: Remove duplicate sequences from each individual dataset. Sequences that appear in multiple distinct input files will remain
- `parallel_process = [True/False]`: Speed up biophysical property matrix generation and distance calculations by utilizing all available cores for the calculations. Will slow other computer operations significantly, and has not been tested on all machines (i.e. Windows OS), which could lead to unexpected issues

- **Note 10:** The choice of alignment will affect all downstream analysis, and can be a place to alter results in re-analysis of a given dataset. Typically, the center alignment scheme is most useful for Abs/TCRs, as the center of CDR loops are often the areas that contact pMHC [9]. In central alignment, information near the N- and C-termini are averaged out (due to differences in sequence length). However, in Abs/TCRs, this information can be recaptured in the V- and J-gene usage. Due to the lack of an analogous information source for peptide N- and C-termini, the bulge is more appropriate. Here, the N- and C-terminal residues, which contain important anchors, are aligned. The remaining central residues then adopt the center alignment, again indicative of the residues most likely to be contacted by TCR [9]. Various values of `bulge pad` should be tested depending on the input dataset.
- **Note 11:** By default, the AIMS analysis is run with 62 distinct biophysical properties that are used to generate position-sensitive metrics and the high-dimensional biophysical property matrix (see Boughter and Meier-Schellersheim [9] for full list). Each of these properties is normalized such that the L1 norm of the 20 values for each amino acid is equal to one. This prevents any one property from dominating in the downstream analyses simply due to higher numeric values for each amino acid property quantification. Users may want to forgo this normalization if they are interested in specific property values (say a precise isoelectric point or hydrophobicity value).
- **Note 12:** This renormalize option is for honing the analysis to focus on the regions of highest sequence diversity in the dataset. This is most noticeably important in, for instance, large TCR datasets. In these cases, V- and J-gene usage dominate the clustering of sequences. Re-weighting by sequence diversity (as quantified by Shannon entropy, discussed in **Note 22**) helps to alleviate this issue. Depending on the application, this may be similarly important in peptide analysis (e.g. peptides eluted from a single MHC should not need this re-normalization, peptides from large datasets may require it).
- **Note 13:** In dimensionality reduction, users may want to use either a `full, parse, or avg` form of the biophysical property matrix. You may want to use a given form of the matrix if:

- `full`: A matrix of size # sequences x # positions in encodings x 62 properties. The full option could be useful if you think that there is a small percentage of your dataset that are massive outliers from otherwise highly correlated features. i.e. your data is being thrown out with the noise of the highly correlated vectors
- `parse`: A parsed version of the full matrix with highly-correlated vectors removed. This is the default in AIMS. Retains most of the important information while reducing matrix size to speed up calculations
- `avg`: A simple per-sequence average of each biophysical property. While potentially over-simplifying much of the data, it can be incredibly useful for testing broad hypotheses. Has been previously shown to strongly discriminate between peptide-presenting and lipid-presenting MHC and MHC-like molecules [12]

- **Note 14:** The choice of dimensionality reduction algorithm will strongly affect the downstream cluster membership. Choices include principal component analysis (‘`PCA`’), t-distributed stochastic neighbor embedding (‘`TSNE`’), or uniform manifold approximation and projection (‘`UMAP`’). Each algorithm has its own strengths and weaknesses:

- `PCA`: As a linear and deterministic algorithm, PCA is the most interpretable and reproducible approach. The principal components identified give back the dimensions of the data with the highest variance in the dataset. So, in a way, you can consider the most distal sequences in the projected PC space to be the most “biophysically distinct” within your dataset
- `UMAP`: UMAP uses nonlinear, stochastic methods to optimize the projection, with the overall goal of preserving the general structure of the sequences in the highdimensional space. This means that no information regarding the “key components” or biophysical properties which determine the “closeness” of certain receptors can be gleaned from this projection. Further, it means that unless users set a specific `random_state` in their calling of the UMAP algorithm, there is no guarantee that their projection will be reproducible. However, it typically does a great job of creating tight clusters of biophysically similar sequences
- `TSNE`: T-distributed Stochastic Neighbor Embedding (t-SNE) is largely similar to UMAP, in that is looking for a lower-dimensional projection of the data which preserves the general structure of the high-dimensional data. However, use of t-SNE is largely deprecated in AIMS, as it performs much worse than UMAP in all applications tested so far

- **Note 15:** Choosing the “proper” clustering algorithm will depend on the input datasets and the hypotheses being tested by the user. Currently, the options include (hierarchical) density-based spatial clustering of applications with noise (‘`HDBSCAN`’ or ‘`DBSCAN`’), k-means clustering (‘`KMEAN`’), or ordering points to identify clustering structure (‘`OPTICS`’):

- `DBSCAN/HDBSCAN`: The (H)DBSCAN algorithm is a density based algorithm, identifying regions of high sequence density surrounded by regions of low sequence density. Due to the extremely high variance in the projected landscape of sequences, what constitutes a “proper” change in density must be user defined (See **Note 16**). HDBSCAN is conceptually similar, but includes an additional step of adding a hierarchical ranking of distances in the projected space using dendrograms
- `KMEAN`: The KMeans algorithm is the simplest of the three used in AIMS, and is the one most appropriate when there exists a strong reason *a priori* for a specific number of clusters. KMeans requires the user to pre-define the number of clusters, and then optimizes the number of points in each cluster based upon distance metrics. Importantly, how this optimization is done in the KMeans algorithm can cause “obvious” clusters to not be properly identified
- `OPTICS`: The OPTICS algorithm is conceptually similar to the DBSCAN algorithm, but with a more user-friendly metric for what constitutes a proper cluster. In the OPTICS algorithm, the user must define the minimum number of sequences that are allowed to be considered a cluster, and the algorithm will go from there in defining clusters based on this minimum number. Each cluster is defined based upon some minimum distance between points, satisfying the minimum cluster number

Note, in most AIMS applications, the “proper” number of clusters will not be obvious *a priori*, so either the OPTICS or DBSCAN algorithm should be used.

- **Note 16:** In choosing the clustering algorithm, users will also need to define hyperparameters for each individual algorithm. The algorithm and associated parameter are as follows:

- `Kmean [NClust]` : `NClust` is simply the number of clusters you would like the KMeans algorithm to generate
- `Optics [min_samples]`: `min_samples` defines the minimum number of samples that can constitute a cluster. Any identified group of sequences that fall below this threshold will be left unclustered
- `DBSCAN [eps]`: `eps` defines the radius of the cluster search space. Due to the different algorithms used in dimensionality reduction, this value can vary widely. Users are encouraged to test multiple `eps` values
- `HDBSCAN [min_cluster_size]`: Identical to the role of `min_samples` as defined in the OPTICS clustering

- **Note 17:** Metadata in AIMS can take two forms, either quantitative or categorical. Categorical metadata is more straightforward to work with, as it provides discrete labels for each individual sequence from which users can selectively isolate in downstream steps. Quantitative metadata, on the other hand, is particularly useful in applications such as single cell RNA sequencing with paired TCR sequencing. In such applications, AIMS cluster plots can be overlaid with counts of a given gene to search for genetic patterns potentially associated with TCR biophysical properties. While incredibly informative, it does become more difficult to select precise sub-groups of data without careful partitioning.
- **Note 18:** Both the clustering and the biophysical property analysis steps in AIMS are where users are most encouraged to test many different software options and subsets of sequences when analyzing the data. How users go about this is very much dataset dependent. If you are starting with a strong hypothesis comparing two or more distinct datasets (e.g. treated v. untreated samples), most analysis should be done on the entire input datasets. If, however, a single dataset or 3+ datasets are loaded into AIMS, typically it is recommended users try to group sequences in an unbiased manner using the dimensionality reduction and clustering algorithms. Analysis can then be carried out on large clusters of sequences.
- **Note 19:** When re-visualizing the selected data subsets (either all identified clusters or all metadata labels), there are two key features that can be toggled. First, lines separating the individual clusters or labels can be added or removed from the plots. In some cases, especially when the DBSCAN or OPTICS algorithms identify a very large number of clusters, this is important for proper visualization of the data. Second, users can optionally toggle a biophysical property view of the groups of data. This allows users to identify populations with striking biophysical features for a closer comparison later in the analysis.
- **Note 20:** Every quantitative comparison between datasets in AIMS comes with the option to calculate bootstrapped standard deviations for the figures and a two-sided nonparametric Studentized bootstrap for significance testing. Both of these calculations can be slow, especially for large datasets. As such, it is recommended that users first explore their datasets without bootstrapping or significance testing, then go back and re-analyze with these additional calculations by simply toggling `bootstrap` and `test sig` values to `True`.
- **Note 21:** In AIMS, publication-ready figures typically display bootstrapped means, standard deviations, and significance testing. This is necessary, as most statistical tests assume normally distributed data, and the majority of sequence datasets are not normally distributed in biophysical property space. Think, for instance, of charge. There are 4 discrete values (+1, 0, -1, partial charges on His and Cys), and as such the charges at a given position in a sequence cannot be normally distributed. Bootstrapping effectively “normalizes” the data and provides reliable estimates of error and statistical significance.
- **Note 22:** In considering the analysis of large sequence datasets, we can take inspiration from more general treatments of written language, which in the abstract is an arrangement of letters that collectively convey information to the reader. Peptides likewise convey information to the immune system through the pMHC-TCR interaction. One such generalizable analytical tool for the study of language and communication is Information Theory [16], which is fundamentally concerned with quantifying the amount of information contained in sets of discrete communicative units. In AIMS, the most used information theoretic metric is Shannon entropy:

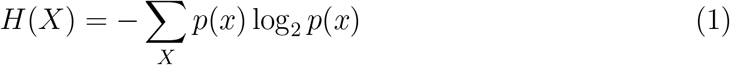

Which is used to quantify the sequence diversity at each encoded position in the AIMS matrix. The minimum entropy is 0 bits (complete amino acid conservation at a given position) while the maximum is 4.32 bits (all 20 amino acids found at a given site with equal frequency). This entropy can then be used to calculate the mutual information, which will be discussed in more detail in **Note 23**. The mutual information provides a more sensitive measurement of the statistical relation in amino acid frequencies at two given positions. It is quantified by first calculating the conditional entropy:

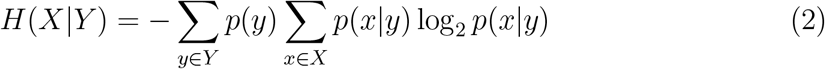

And then subtracting this conditional entropy from the standard entropy at a given position to give the mutual information *I* :

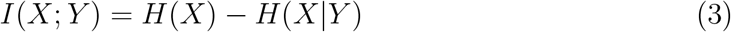

- **Note 23:** Similar to what was mentioned in **Note 20**, significance testing and bootstrapping of mean values and standard deviations is even slower for information theoretic metrics. It is strongly recommended that statistical significance is assessed only once data exploration is complete and final quantifications are initiated.
- **Note 24:** As a first pass, mutual information between two positions in the encoded AIMS matrix can be considered a correlation between the residues at those sites. However, mutual information is more sensitive than correlation, as it excludes highly conserved amino acids, which can be detected using other methods (such as the Shannon entropy plots shown in **Figure 9**). As such, you can consider residue positions with a high mutual information as either directly interacting within a given structure, collectively coordinating interactions with another molecule, or under some other selective pressure which still allows for some level of amino acid diversity. Again, such nuance is important in the analysis of immune molecules due to conserved anchors in peptides or conserved disulfide bridges in TCRs and Abs.
- **Note 25:** The usage of N-gram analysis in AIMS was again inspired by Claude Shannon’s original work on information theory [17]. They are incredibly useful to find patterns that mutual information may not catch, if for example an RK consecutive motif is well conserved in a set of peptides, but that RK does not appear at the same sequence position. Generally, users should not go above tri-gram analysis, as in addition to the difficulty of visualization, N-gram frequency decreases drastically at each higher order.
- **Note 26:** For simplicity, the AIMS classification module has been standardized to use linear discriminant analysis, the recommended classifier (see **Note 27** for explanation why). However, users of the Jupyter notebook or AIMS CLI can access a larger suite of classification algorithms through the `do_classy_mda` function and select from machine learning algorithms including `support vector machine, logistic regression, or random forest`, along with a range of options for parsing through the high dimensional matrix and assessing model accuracy.
- **Note 27:** It should be noted that machine learning models have overall performed poorly on immunological datasets. While models can be generated for certain modest features of a given sequence (e.g. likelihood of a peptide to bind to a certain HLA allele, polyreactivity status of an antibody, etc) broader models such as predictions of which TCR-pMHC pair will elicit an immune response generalize poorly [18,19]. If generating and publishing a machine learning model using AIMS, exceptional care should be taken to reduce the likelihood of overfitting. The benefit of using linear discriminant as the classifier is its inherent interpretability. Analysis of the linear weights allows users to identify why certain sequences are classified into a given class, leading to increased transparency that provides a readout to assess the likelihood of model overfitting. The majority of other classification algorithms are so called “black-boxes” preventing users from identifying the source of high classification accuracy.
- **Note 28:** Assuming that users choose to generate a classifier using AIMS after having considered the issues with machine learning discussed in **Note 27**, they will still need to test a range of hyperparameters to create an optimal model. While a full discussion of hyperparameters is outside of the scope of this work, the most important thing to note is that the selected `matSize` should be significantly smaller than the number of sequences in the smallest dataset used in the binary comparison. Otherwise, each feature can be tuned for each individual sequence in this dataset to achieve an artificially high classification accuracy.
- **Note 29:** As a final point, it is important to note that AIMS is constantly changing. While the core functionality should never stray significantly from what is found in this chapter, there may be additional useful features that are added after the publication of this textbook. For the most up-to-date information on AIMS, users should check the documentation website: `aims-doc.readthedocs.io`

## Acknowledgements

This work was supported by the intramural program of the National Institute of Allergy and Infectious Diseases (NIAID), NIH. This is a preprint of the following chapter: Christopher T. Boughter, Utilizing Protein Bioinformatics to Delve Deeper into Immunopeptidomic Datasets, published in *Methods In Molecular Biology Series: Immunoproteomics - 3rd Edition*, edited by Kelly M. Fulton and Susan M. Twine, 2025, *Springer Protocols*. Reproduced with permission of Springer Science+Business Media, LLC, part of Springer Nature. The final authenticated version will be published online, and the DOI will be included here in a later update to this preprint.

